# High density genome scan for selection signatures in French sheep reveals allelic heterogeneity and introgression at adaptive loci

**DOI:** 10.1101/103010

**Authors:** Christina Marie Rochus, Flavie Tortereau, Florence Plisson-Petit, Gwendal Restoux, Carole Moreno, Gwenola Tosser-Klopp, Bertrand Servin

## Abstract

Sheep was one of the first domesticated livestock species in the Anatolia region of contemporary Iran and eventually spread world-wide. Previous studies have shown that French sheep populations likely harbour a large part of European domesticated sheep diversity in a relatively small geographical region, offering a powerful model for the study of adaptation. We studied the diversity of 27 French sheep populations by genotyping 542 individuals for more than 500 000 SNPs. We found that French sheep breeds were divided into two main groups, corresponding to northern and southern origins and additionally we identified admixture events between northern and southern populations. The genetic diversity of domesticated animals results from adaptation of populations to constraints imposed by farmers and environmental conditions. We identified 126 genomic regions likely affected by selection. In many cases, we found evidence for parallel selection events in different genetic backgrounds, most likely for different mutations. Some of these regions harbour genes potentially involved in morphological traits (*SOCS2*, *NCAPG/LCORL*, *MSRB3*), coat colour (*MC1R*) and adaptation to environmental conditions (*ADAMTS9*). Closer inspection of two of these regions clarified their evolutionary history: at the *LCORL/NCAPG* locus we found evidence for introgression of an adaptive allele from a southern population into northern populations and by resequencing *MC1R* in some breeds we confirmed different mutations in this gene are responsible for the same phenotypic trait. Our study illustrates how dense genetic data in multiple populations allows the deciphering of evolutionary history of populations and of their adaptive mutations.

## Introduction

Sheep, one of the first domesticated species, originated from a population of *Ovis orientalis* in the fertile crescent in contemporary Iran (Dobney and Larson, 2006; Zeder, 2008). From its domestication centre, sheep eventually spread world wide, adapting to a large range of environmental conditions. In agriculture, sheep are raised for meat, wool and milk products, with commercial breeds usually specialized for one type of production. This history has shaped sheep genetic diversity and makes it an interesting model species to study the evolution of populations under adaptive constraints (Andersson, 2012).

In a recent study, the International Sheep Genomics Consortium (ISGC) (Kijas *et al.*, 2012) compiled a dataset of DNA samples from 74 populations of sheep from around the world genotyped for approximately 50 000 SNP markers: the SheepHapMap dataset. Analyses of this dataset showed both clear geographical structure of breeds and strong admixture among breeds. In Western Europe, geographical structure was clear with several population groups identified that corresponded to their area of origin: Italian, Iberian, Swiss, British and Northern European groups (Kijas *et al.*, 2012). Only one French sheep population, the Lacaune breed, was included in this study. It clustered in the Iberian group, consistent with its origin from the south-western region of France. Besides the Lacaune breed, France harbours many other different sheep breeds across the country and because of its geographical position, France borders many of the geographic areas associated with western European sheep population groups of the SheepHapMap dataset. The study of French sheep diversity should therefore provide valuable insights on the establishment of genetic structure in European sheep, and in particular inform us on the history of sheep husbandry in Europe. French sheep diversity was recently studied with the genotyping of 21 microsatellites in sheep from 49 breeds (Leroy *et al.*, 2015). This study confirmed that French sheep breeds show a structure consistent with influences from both Northern Europe, specifically British breeds, and southern European breeds. Hence, French sheep populations are likely to harbour a large part of European sheep diversity in a relatively small geographical region, offering a powerful model for the study of adaptation.

Several studies have identified signatures of selection in sheep: signatures associated with pigmentation (Garcia-Gamez *et al.*, 2011), fat deposition (Moradi *et al.*, 2012), milk yield (Moioli *et al.*, 2013), adaptations to climate (Lv *et al.*, 2014) and resistance to parasites (McRae *et al.*, 2014). There have also been studies looking at a number of breeds: a study which included only American breeds of sheep (Zhang *et al.*, 2013) and two studies including sheep populations from around the world (Kijas *et al.*, 2012; Fariello *et al.*, 2014). These worldwide studies, which both used data from the ISGC SheepHapMap data set, detected signatures containing genes or QTLs associated with pigmentation, morphology, muscling, milk production and wool quality. Studying data sets with many different breeds also allows the study of diversity in selection, the different mutations and genes selected on to reach the same phenotype (ie. colour, muscling etc.). All of these studies however, relied on medium density genotyping using about 50 000 SNP markers. This limits the precision of localization of candidate genes underlying selection signatures as well as the identification of potential haplotypes carrying causal mutations.

The present study aims to enhance our understanding of adaptation in sheep by providing and analyzing a large data set of high density genotyping (around 600 000 SNP markers) in 542 samples from 27 French sheep populations. Our analyses of the genetic structure in this dataset identify clearly two main origins of breeds and show how these breeds complement the SheepHapMap dataset. Then, in what is the first selection scan based on high-density genotyping in sheep, our results show clear evidence for adaptive introgression between groups along with allelic heterogeneity at some adaptive loci. Finally, we confirm this heterogeneity by re-sequencing a known coat colour gene (*MC1R*) in three populations presenting evidence for selection at this locus, and pinpoint a set of new potential causal mutations for coat colour phenotypes in sheep.

## Materials and Methods

### French sheep data

A total of 691 French sheep were genotyped for 653 305 autosomal SNPs (Ovine Infinium^®^ HD SNP BeadChip) with 27 sheep breeds being represented in this study. Breed names and their short forms are listed in Table 1. In each breed genotyped animals were chosen so as to be as unrelated as possible based on pedigree records. DNA samples from this study came from various French flocks. Blood sampling was not performed specifically for this study. Blood samples were taken from commercial farms. Animals did not belong to any experimental design but were sampled by veterinarians and/or under veterinarian supervision for routine veterinary care. A test for Hardy Weinberg Equilibrium (HWE) was calculated for each SNP within each breed using PLINK (Purcell and Chang, 2015) and SNPs not is HWE (False discovery rate (FDR) 5% calculated with R package qvalue (Storey, 2015)) in one or more breeds were removed. For each breed, a genomic relationship matrix was computed (Yang *et al.*, 2011). The resulting distribution of kinship coefficients was modelled as a mixture of normal distribution, with the major component of the mixture representing pairs of unrelated individuals. Pairs of related individuals were identified as those that did not belong to this component (FDR < 5%). Finally, one individual for each pair was removed until no related individuals remained. All further analyses were performed on this set of unrelated individuals. SNPs with a minor allele frequency (MAF) = 0, SNPs with a missing call rate > 0.01 and sheep with individual missing call rate > 0.05 were removed.

**Table 1:** Sheep population groups and breeds

| Origin | Breed | Abbreviation |
| --- | --- | --- |
| Mouflon | European Mouflon | EUR |
|  | Asian Mouflon | IRN |
| Northern Europe | Berrichon du Cher | BER |
|  | Charollais | CHA |
|  | Charmoise | CHR |
|  | Île de France | IDF |
|  | Ouessant (Ushant) | OUE |
|  | Romane | RMN |
|  | Romanov | ROM |
|  | Roussin de la Hague | ROU |
|  | Rouge de l'Ouest | RWE |
|  | Suffolk | SUF |
|  | Texel | TEX |
|  | Vendéen | VEN |
| Southern Europe | Blanche du Massif Central | BMC |
|  | Causse du Lot | CDL |
|  | Corse | COR |
|  | Dairy Lacaune | LAC |
|  | Meat Lacaune | LAM |
|  | Limousine | LIM |
|  | Mérinos d'Arles | MER |
|  | Mérinos de Rambouillet | RAM |
|  | Mourerous | MOU |
|  | Manech tête rousse | MTR |
|  | Noire du Velay | NVE |
|  | Préalpes du Sud | PAS |
|  | Rava | RAV |
|  | Tarasconnaise | TAR |

### Population structure of French sheep

Three methods were used to study population structure in French sheep from SNP genotypes: A principal components analysis to visualize patterns in relationships between individuals using PLINK (Purcell and Chang, 2015); A model based approach to estimate individual ancestry coefficients, using the software sNMF (Frichot *et al.*, 2014); A model based approach to infer populations splits and mixtures using the software treemix (Pickrell and Pritchard, 2012). For sNMF, the optimal number of ancestral populations (K) was the one that had the lowest cross entropy criterion (Supplementary Figure 1) (Frichot *et al.*, 2014). For treemix, only the 498 651 SNPs genotyped in both Asian Mouflon and French sheep were included. The number of migration events included in the population tree was chosen based on the comparison of the fraction of the variance in relatedness between populations that is accounted for by the model, as explained in the software documentation (Supplementary Figure 2).

### Population structure of European sheep

To study French sheep population structure within other European breeds, a subset of breeds from the Sheep HapMap data set were included in population structure analyses; both model based approaches, estimation of individual ancestry coefficients and inferred population splits and mixtures. The SheepHapMap individuals included in this study were the same European sheep used by Fariello et al. (Fariello *et al.*, 2014) including breeds from; Central Europe; South Western Europe; North Europe; and Italy. Breeds were removed if they had a sample size < 20, or were a result of recent admixture or had recently experienced a severe bottleneck (Fariello *et al.*, 2014). Shared markers between the French sheep breeds and the SheepHapMap subset of breeds, 39 976 SNPs, were used to estimate individual ancestry coefficients using sNMF (Frichot *et al.*, 2014). The same Iranian Asian Mouflon sheep were included in the analysis to root the population tree when estimating population splits and mixtures using treemix (Pickrell and Pritchard, 2012) where allele frequencies from 30 Iranian Asian Mouflon sheep from the NEXTGEN project were included to root the population tree. For the combination of our dataset with the SheepHapMap dataset, a total of 38 596 SNPs common to the French sheep, SheepHapMap subset and Asian Mouflon sheep were used in the analyses.

### FLK and hapFLK genome scans

FLK (Bonhomme *et al.*, 2010) and hapFLK (Fariello *et al.*, 2013) tests were used to detect signatures of selection. Both tests are aimed at identifying regions of outlying differentiation beteen population while accounting for their hierarchical structure. The evolutionary model underlying FLK and hapFLK assumes that SNPs were polymorphic in an ancestral population. In order to consider only SNPs matching this hypothesis, SNPs with estimated ancestral minor allele frequency < 5% were removed, leaving 465 2351 SNPs for use in the analyses. For the FLK analysis, significant SNPs within 1 Mbp from each other where grouped into a common selection signature (FDR 1%). Candidate genes were identified in regions containing 10 or more significant SNPs. For hapFLK, a number of haplotype clusters has to be specified in order to fit the Scheet and Stephens model (Scheet and Stephens, 2006). This number was set to 30, from results obtained using the cross validation procedure included in the fastPHASE software. P-values for hapFLK were obtained by fitting a chi-squared distribution to the empirical distribution as explained in ref. (Boitard *et al.*, 2016) using the script available on the hapflk software web page. For the hapFLK analysis, regions were constructed from significant SNPs (FDR 5%) and grouped together if they were 1 Mbp or less apart. The FLK analysis was run with all breeds and northern and southern breeds separately and these results were compared to determine the regions missed when northern and southern breeds are run separately. The hapFLK statistics were computed for northern and southern groups separately. Candidate genes were searched for in regions with five or more significant SNPs. Regions from both FLK and hapFLK results had all protein coding genes present extracted. The SNP with the lowest p-value in each region is referred to as the best SNP in Supplementary Table 1. The distance from each gene to the best SNP was determined by the distance of the midpoint of the gene to the best SNP and then genes were than ranked with the closest gene labelled as the best gene. Of all results, nine regions were selected for further analysis based on the candidate genes located within the regions. Local trees were constructed by recomputing Reynold’s distances between populations in a region and re-estimating branch length of the whole genome tree from local Reynold’s distances as in (Fariello *et al.*, 2013). Local trees from single SNPs and allele frequencies in FLK results were evaluated to determine breeds selected on and for hapFLK results local trees from single SNPs and haplotypes, allele frequencies and haplotype cluster frequencies.

### Additional polymorphism discovery and large-scale genotyping at MC1R locus

The *MC1R* locus (OAR14:14 228 283-14 235 506 on OARv3.1 assembly) was amplified as overlapping PCR fragments using appropriate PCR conditions regarding the expected length of the product: either conventional amplification using goTaq polymerasae (Promega) or Long-range PCR amplifications using the Long PCR Enzyme Mix provided by Fermentas (http://www.fermentas.de) was performed, using 50 ng of genomic DNA as a template and the manufacturer’s protocol. After treatment with 0.5 U of Tsap (Promega) and 10 U of exonucleaseI (Biolabs), 10 to 90 ng of PCR product were used for sequencing with either internal primers or PCR primers used for the amplification. Sequencing reaction was carried out via the BigDye^®^ Terminator v3.1 Cycle Sequencing Kit (http://www.appliedbiosystems.com). The primers are listed in Supplementary Table 2. Sequences from three animals from each breed exhibiting a selection signature were aligned against scaffold 00839 of the *Ovis aries musimon* assembly (GenBank accession HG925721.1), which we found of better quality than the OAR3.1 reference genome, using CLC software and allowed the detection of polymorphisms. The discovered polymorphisms were genotyped for all the animals of the relevant breeds by sequencing purified PCR products with internal primers using the same protocol. The sheep genome browser from EBI (www.ensembl.org) was used to determine conserved regions through 39 eutherian mammals and calculate GERP scores.

## Results and Discussion

### French sheep divide into two main geographical groups

Samples from 27 French populations were genotyped using a high density SNP array for 527 823 SNP markers after quality control. The populations chosen represent the majority of commercial breeds present in France but also included some breeds maintained for conservation purposes. We inferred population structure in French sheep using three different approaches: a principal component analysis (PCA) based on the genotype matrix, an unsupervised clustering approach based on Hardy Weinberg Equilibrium within clusters, similar to the STRUCTURE model (Pritchard *et al.*, 2000) but with better computational properties on large samples (Frichot *et al.*, 2014) and a model-based approach aimed at reconstructing a population tree with possible admixture events (Pickrell and Pritchard, 2012).

To get insight into the domestication history of the breeds, we included two outgroup populations in our analyses. Allele frequencies in the Asian Mouflon (the ancestor of domesticated sheep (Nadler *et al.*, 1973; Bunch *et al.*, 1976)) were obtained from the NextGen project and were used to root the population tree in the maximum likelihood tree analyses. We also included two genotyped samples of European Mouflon from Corsica in the principal component, individual ancestry coefficient and maximum likelihood tree analyses. The placement of the European Mouflon sheep on the maximum likelihood tree analyses (Figure 4D), after the Asian Mouflon sheep but before any other domesticated breeds, was consistent with the known origin of this population: European Mouflon sheep were domesticated sheep that became feral early after their arrival to Europe and can still be found on Corsica and Sardinia islands in the Mediterranean today (Hermans, 1996). Because the European Mouflon sheep appeared on the tree before any of the French sheep, we used these two animals in selective sweep studies as an outgroup to root the population tree.

**Figure 4.**
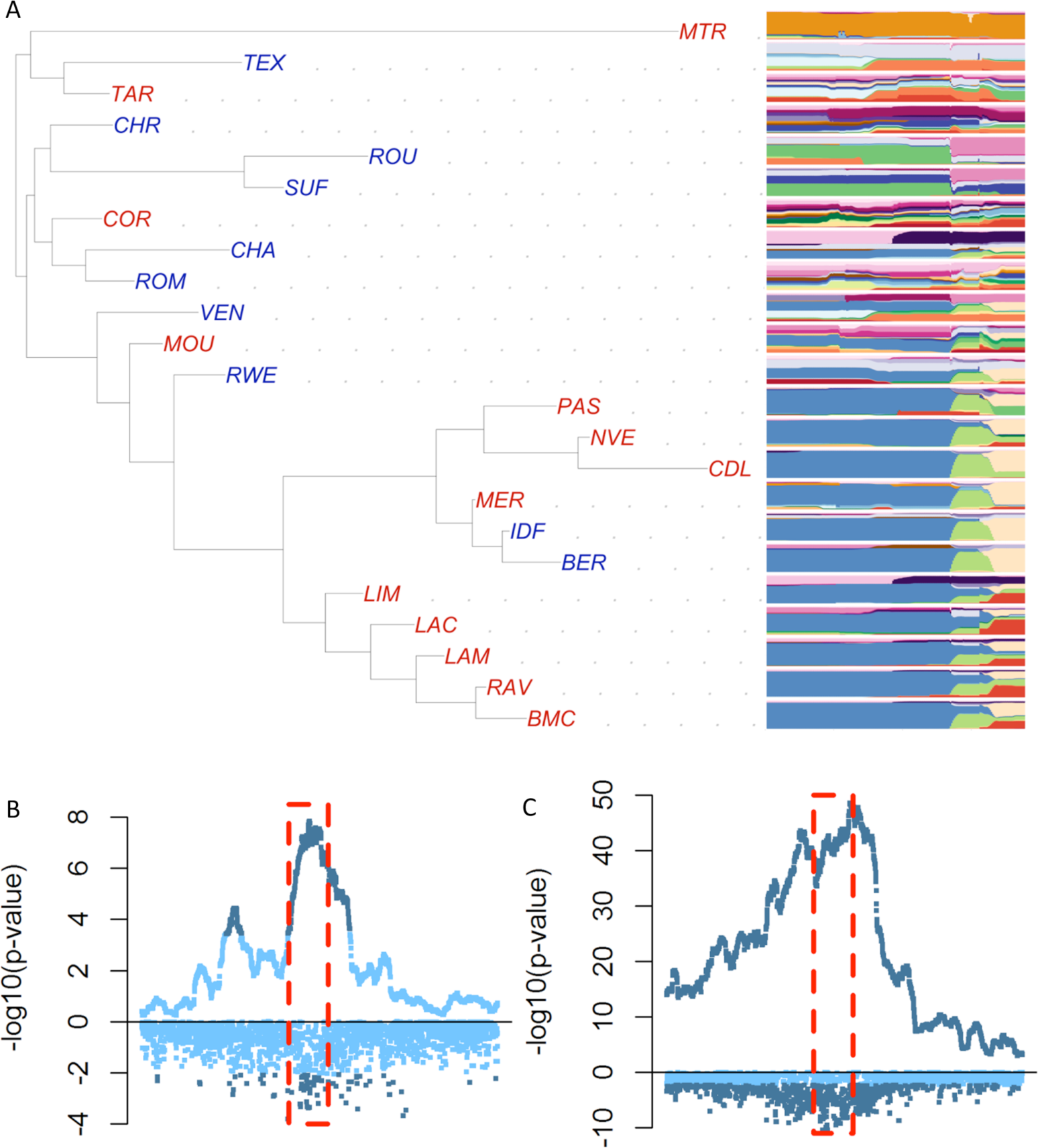
**Selection signature around genes ABCG2,NCAPG and LCORL on chromosome 6.** (A) Population tree (left) constructed in the region under selection with haplotype cluster plots shown for each breed (right). hapFLK and FLK p-values for the selection signature in the northern (B) and southern (C) populations.

In domesticated samples, the three approaches showed a clear structure (Figure 4) in two main groups of breeds plus two highly differentiated breeds, the Ouessant (also called Ushant) sheep and the Mérinos de Rambouillet. The Ouessant sheep breed has historically been isolated and consequently has accumulated substantial drift. Its position in the tree is consistent with it having a distant origin from the other breeds in the data set. The Ouessant breed possibly originated from a first wave of colonization after domestication, as has been shown for other ancient European breeds of sheep such as the Soay sheep (Chessa *et al.*, 2009). However, its positioning in the population tree might not be very reliable due its long branch length. The other highly inbred breed is the Mérinos de Rambouillet. This breed consists of a single flock of animals that has been raised without the introduction of additional animals since 1786. Although it has a very long branch in the population tree, it clusters with the other Merino population in the dataset, the Mérinos d’Arles.

Apart from the Ouessant and Mérinos de Rambouillet, the populations were divided into two main groups, corresponding to northern and southern origins (Figure 4). The two groups present contrasting structure. In the northern group, breeds were clearly separated on the PCA analysis (Supplementary Figure 3), individuals had clear cluster assignment (Figure 4B) and tended to have longer branch length in the population tree (Figure 4D), indicating higher drift and differentiation between breeds. In contrast, breeds of southern origin show less differentiation, with PCA clusters being closer, individual clustering subtler and shorter branch length in the population tree. This pattern is particularly clear for breeds originating in the Massif Central (Blanche du Massif Central, both Lacaune populations, and Noire du Velay), but is also true of breeds more geographically distant such as the Manech Tête Rousse and the Préalpes du Sud. The division between northern and southern French breeds in population structure analysis is possible evidence of the effects of routes of domestication on modern sheep breeds and because of longer established formal breeding programs in northern Europe.

The population tree that best explained genetic relationships between breeds had four migration events (Figure 4D). The migration edge with the most weight was from the Romanov to the Romane which was on the same branch as the Berrichon du Cher. This migration is explained by the fact that the Romane breed was created in France in the 1960s by crossing Romanov and Berrichon du Cher animals. The migration edge from the Mérinos de Rambouillet branch to the branch of both Île de France and Berrichon du Cher breeds is likely representative of the crossing of these breeds with Merinos in the nineteenth century to improve wool quality (Porter, 2002). The migration edge from the Roussin de la Hague branch to the branch with Texel, Rouge de l’Ouest, Île de France and Berrichon du Cher breeds could come from the use of British breeds to crossbreed in the eighteenth century to improve meat production (Porter, 2002). However this is difficult to confirm because the main British breeds involved, the Leicester Longwool and Southdown, are not part of our dataset nor of the SheepHapMap dataset. Finally, the migration branch from the Île de France to Mourerous breed was an unexpected result and might be due to some purported recent crossing of the Mourerous with northern breeds to improve meat production.

### French sheep populations and the meeting of European sheep domestication routes

To place the population structure of French sheep into the context of European population diversity, population tree with mixture events and individual ancestry coefficients were estimated using European sheep breeds from the SheepHapMap project and included the as the French data set downscaled to medium density SNP information. The maximum likelihood population tree with no migration events (Figure 4) showed that French sheep complement the SheepHapMap dataset in the European sheep population tree. Most of the breeds in our dataset added to the global diversity, as they tend to root in internal branches of the population tree, either within northern or southern breeds (*e.g*. the Berrichon du Cher/Île de France and Causse du Lot/Rava/Limousine respectively) or at the basis of the population tree (*e.g*. the Ouessant or Romanov breeds). The other French breeds branched in population groups that are already present in the SheepHapMap: the Corsican breed is closely related to Italian breeds, and the Sardinian Ancestral Black in particular; the other cases corresponded to breeds that are also found in other countries and the population tree reflects the diversity within these breeds between countries (examples include the French and Irish Suffolk populations, the German, Scottish, New Zealand and French Texel sheep populations, and the Australian Merino, Mérino d’Arles and Mérino de Rambouillet breeds).

As when considering French breeds only, the tree obtained from the inclusion of the SheepHapMap breeds separated into: southern breeds, which included Spanish and Italian breeds; and northern breeds. Here also, the northern breeds tended to have longer branch lengths and more drift than southern breeds. When estimating ancestral populations at K=3 there was differentiation between northern European and southern European sheep breeds similar to the analysis with only French breeds at four ancestral populations (K=4) (Figure 4B). At K=20 (the K value with the lowest cross-validation error rate, Supplementary Figure 4), breeds such as the Texel showed large variation both within the breed and within regional populations such as the Scottish Texel).

### Selection signatures are mostly detected within geographical groups

To detect signatures of selection we used two methods, FLK and hapFLK, which are both based on outlying differentiation between populations (a single marker approach and its haplotype based extension) that accounts for the hierarchical structure between populations (Bonhomme *et al.*, 2010; Fariello *et al.*, 2013). While hapFLK methods are robust to bottlenecks and moderate admixture, both of these can affect power for detecting signatures of selection (Fariello *et al.*, 2014). We therefore did not consider four of the breeds for our selection scan: three breeds, the Ouessant, Mérinos de Rambouillet and Berrichon du Cher, that have experienced severe bottlenecks, corresponding to very long branch lengths in the population tree (Figure 4D); and the Romane breed, as it is a recent composite of two breeds.

We detected selective sweeps with all breeds taken together and separately in northern and southern sheep breeds using both FLK and hapFLK (results in Supplementary Table 1). When considering all breeds together, we detected 50 selection signatures. Three of these signatures were only found when analyzing all the breeds together, while the other 47 were also found in the within group analyses. These three specific signatures were on chromosome 3, between 129.0 Mb and 130.7 Mb; on chromosome 8, between 80.6 Mb and 83.1 Mb; and on chromosome 24 between 9.9 and 10.6 Mb (see File Supplementary Table 2 for detailed figures of all selection signatures).

The signature on chromosome 3 harbours the *SOCS2* gene recently shown to carry a mutation with adverse pleiotropic effects, positive for growth and negative for mastitis (Rupp *et al.*, 2015). The pattern of haplotype diversity showed that a haplotype cluster segregates at moderate frequency in many northern breeds (light yellow on page 18 from Supplementary Table 3), that can only be detected as outlying by including southern breeds. Moderate allele frequencies at this common haplotype in many northern breed could be due to balancing selection on adverse pleiotropic effects at *SOCS2*, although it is essentially impossible to test this hypothesis from SNP array data alone. Analysis of the region on chromosome 8 is consistent with selection in Île de France, although the selected haplotype cluster has not reached complete fixation in this breed. It was also found at low frequency in the Texel and Rouge de l’Ouest populations, closely related to Île de France. Hence, when analyzing only the northern breeds, this signal can still be consistent with drift alone. However, as none of the southern breeds harboured this haplotype, haplotype diversity patterns in northern breed become unlikely due to drift alone. Finally, the region on chromosome 24 exhibited a very low haplotype diversity in most northern and southern breeds, with a similar haplotype segregating at high frequencies in many breeds (*i.e*. low differentiation due to general selection on the same haplotype). Only a few northern breeds had some more elevated haplotype diversity, which allowed detecting selection when included with the southern breed. Interestingly, the selection signature in this region looked quite widespread for the same haplotype in most of the breeds in the dataset.

As only three signatures were specific to analyzing all breeds together, we then focused our inference on the analyses performed in each group separately. We had more power to detect signatures of selection when separating breeds into the two groups: for FLK and hapFLK we detected 61 and 26 regions respectively for northern sheep populations and 65 and 42 regions for southern sheep populations. As shown above, southern French sheep are not highly differentiated from each other and have experienced relatively low drift compared to northern ones. Because of this we had more power in southern than in northern populations which could explain the detection of more regions under selection and much smaller p-values in the former than in the latter (Figure 4).

With the high density SNP information selective sweep regions were on average smaller than those detected by Fariello *et al.* (Fariello *et al.*, 2014) using medium density (50K) SNP information. While we acknowledge that it is difficult to make comparisons with Fariello *et al.* (Fariello *et al.*, 2014) because that study included different sheep populations, regions detected with the 50K SNP array were 2.8 times larger than those that were detected with the 600K array (Supplementary Figure 5). For those regions detected in both studies, the regions detected with 50K were 1.7 times the size of those detected with 600K SNPs. The higher resolution obtained can come from the fact that we get a better haplotype diversity description with higher SNP density and are able to reduce the size of the candidate region by analyzing more breeds showing more recombinant haplotypes.

### Overview of candidate genes found in selection signatures

Based on the FLK and hapFLK results, we defined selection signatures as intervals where nearby SNPs were significant (see Materials and Methods). We then listed all the genes in the regions, and in some cases could pinpoint a list of likely candidate genes, based on their distance to the most significant SNP and on previous literature on their functions or selective status. These candidates include genes for coat colour (*ASIP*, *MC1R*, *TYRP1*, *MITF*, *EDN3*, and *BNC2*), stature and morphology (*NPR2*, *MSTN* (*GDF-8*), *LCORL* and *NCAPG*, *ALX4* and *EXT2*, *PALLD*), milk production (*ABCG2*), horns (*RXFP2*) and a region where a wool quality QTL has been found (Kijas *et al.*, 2012; Fariello *et al.*, 2013). Other candidate genes that we identified included *SOCS2*, associated with growth and mammary gland development (Rupp *et al.*, 2015), *OXCT1*, associated with milk fatty acid traits in dairy cattle (Li *et al.*, 2014), EBF2, involved in brown fat fate and function in mice (Rajakumari *et al.*, 2013), *ADAMTS9*, under selection in Tibetan pigs and boars living at high and moderate altitudes and *MSRB3*, associated with floppy ear position in dogs (Boyko *et al.*, 2010). We further discuss some of these selection signatures with candidate genes that have had their function in mammals discussed in peer-reviewed literature and are no further than 150 kbp from the SNP with the smallest p value for FLK or hapFLK tests in the region.

### Origin of the causal mutation in MSTN

In the case of *MSTN*, the causal mutation is a SNP that is part of the HD SNP array used in our study (rs408469734). We found that it was the SNP showing the most significant FLK signal of the corresponding selection signature, and local population trees highlighted the Texel breed as being the selected population (page 14 of Supplementary Table 3). As it is the causative mutation, our dataset allows us to highlight the evolutionary history of this particular mutation: we found this SNP segregating in another meat breed, the Rouge de l’Ouest where it is carried by a similar but smaller haplotype than in the Texel. To evaluate the spread of the signal due to the selection in Texel, we conducted a new FLK analysis on chromosome 2 with all animals except the Texel sheep. Selection signatures found in this analysis are shown in Supplementary Table 4. In contrast to results from FLK using all animals there were five fewer regions detected, spanning together from 109.0 to 122.3 Mbp, surrounding the *MSTN* gene. All the other selection signatures on chromosome 2 were still found. Therefore the Texel selected haplotype is very long, likely stretching from 109.0 to 122.3 Mbp. This shows a very strong and recent selective sweep. Interestingly we still have evidence for selection in this new analysis in the Rouge de l’Ouest, but not associated to fixation of the allele, the signal ranging from 117.9 to 118.8 Mbp. We thus concluded that both breeds share a common ancestral population where this SNP was segregating and that this mutation was not introduced into the Rouge de l’Ouest via introgression from the Texel as only part of the Texel haplotype is seen in the Rouge de l’Ouest population.

### Many selection signatures exhibit allelic heterogeneity

We define allelic heterogeneity as the presence of multiple selected alleles, or haplotypes, at the same genomic location. In our analyses, we identified allelic heterogeneity in selection signatures when we had evidence for (i) more than one breed affected by selection in the same region and (ii) different haplotypes having arisen to high frequency in the selected breeds. We found many examples of allelic heterogeneity in selection signatures, including regions containing genes shown to be involved in agronomically important traits such as *ADAMST9, MSRB3, SOCS2, RXFP2, LCORL* and *MC1R*.

Regions under selection harbouring *ADAMTS9* (*A disintegrin-like and metalloprotease (reprolysin type) with thrombospondin type 1 motif 9*) (page 45 of Supplementary Table 3) and *MSRB3* (*Methionine sulfoxide reductase B3*) (page 19 of Supplementary Table 3) are simple examples of one region with more than one haplotype under selection. *ADAMTS9* has been shown to have been under selection in pigs and wild boars living at high altitudes in Tibet in two studies (Li *et al.*, 2013; Dong *et al.*, 2014). The first study was a comparison of Tibetan wild boars with domesticated southern Chinese pigs living at low altitude (Li *et al.*,2013) while the other was a comparison of domesticated southern Chinese pig breeds which lived at high, moderate and low altitudes (Dong *et al.*, 2014). In mice, the *ADAMTS9* protease has been shown to be involved both in cardiovascular development and homeostasis (Kern *et al.*, 2010). In our study, selection signatures were detected in the region of *ADAMTS9* on chromosome 19 and the SNP with the lowest p-value for FLK analysis for all breeds was located within the gene. Two breeds were found to be under selection for this region: the Romanov and the Causse du Lot breeds. The Causse du Lot breed is a hardy breed traditionally raised in plateaus of southern France and the Romanov breed is a European north short tailed sheep breed originating in Russia and hardy in cold temperatures. This suggests that *ADAMTS9* may have a role in the ability of hardy breeds to survive in colder temperatures.

*MSRB3* is in a region that has been associated with ear position (floppiness) in dogs (Jones *et al.*, 2008; Boyko *et al.*, 2010; Vaysse *et al.*, 2011) and ear size in pigs (Zhang *et al.*, 2014; Zhang *et al.*, 2015). In a study of selection signatures in Chinese sheep, researchers detected selection in the breed with the largest ears in their study (Wei *et al.*, 2015). In our study, there was a hard sweep in Suffolk in this region and a few other haplotypes at high frequency in the Romanov and Charolais breeds. The breed definition for Suffolk sheep in France from the national organization for selection (GEODE) describes ears as long, thin and facing in a downwards direction (Organisme de Sélection Génétique Ovine et Développement (GEODE) 2013). Hence, our results suggest that mutations in *MSRB3* could cause this breed’s phenotype.

The *Relaxin-like receptor 2* (*RXFP2*) gene has been associated with the presence/absence of horns as well as horn development in different breeds of sheep and is therefore already known for its heterogeneity. In Soay sheep, horn type (horns or scurs) and horn length was associated with the same region on chromosome 10 which was mapped to a single candidate gene, *RXFP2* (Johnston *et al.*, 2011). Later, a single SNP was found to be highly predictive for horns (dominant when inherited maternally) in Australian Merino sheep on chromosome 10 (Dominik *et al.*, 2012). A synonymous mutation, p. P375 (c.1125A > G) in Tan and Suffolk sheep was found to be associated with the appearance or absence of horns (Wang *et al.*, 2014). When comparing horned and polled animals from seven Swiss breeds of sheep a 1833 bp insertion in a 4 kb region at the 3 prime end of *RXFP2* was found only in polled animals (Wiedemar and Drogemuller, 2015). Finally, the *RXPF2* region was detected as a signature of selection in different population groups of the SheepHapMap dataset (Kijas *et al.*, 2012; Fariello *et al.*, 2014). The haplotype cluster frequency plots from those breeds under selection highlighted the complexity of selection in this region. The signal in our study was detected in all (page 28 of Supplementary Table 3) and separate analyses of northern and southern sheep with the two most significant SNPs in the southern analyses within *RXFP2* (both intronic mutations) while the signal from hapflk results of northern sheep peaked 89 Kb before the gene. None of the northern breeds included in our dataset are horned, and the reason for detecting a signature here is because different haplotypes have been selected on within this region. When we looked at the region extending 100 kbp on either side of *RXFP2*, while the SNP frequencies within the gene were similar among northern sheep and other polled southern sheep, in the 100 kbp before the gene there were four distinct haplotypes in polled French sheep. This finding demonstrates that it is likely that multiple ancient mutations are affecting polled phenotypes rather than a case of a single mutation being introgressed into all polled breeds.

While different haplotypes under selection don’t necessarily mean multiple functional mutations, we found a possible example of this in the region under selection containing the *Suppressor of cytokine signalling 2* (*SOCS2*) gene (page 18 of Supplementary Table 3). *SOCS2* is thought to play roles in metabolism, somatic growth, bone formation, the central nervous system, response to infections and mammary gland development although the main target of *SOCS2* is believed to be *GH/IGF-1* which is important for somatic growth (Rico-Bautista *et al.*, 2006). In a grand-daughter study of 1009 commercial French dairy rams (dairy Lacaune breed) researchers found a QTL associated with somatic cell count on chromosome 3 and a highly associated SNP in *SOCS2* (Rupp *et al.*, 2015). The frequency of this SNP in the studied population was 21.7% (Rupp *et al.*, 2015). In contrast, in our study we detected selective sweeps in this region in mostly northern French breeds raised for meat. Considering both the literature and the results of this study, there is evidence that there is more than one functional mutation found in *SOCS2* in French sheep.

### Example of selection on adaptive introgression

A region on chromosome six showed both allelic heterogeneity and was an example of selection on a genome region introgressed after an admixture event (page 23 of Supplementary Table 3). In a previous scan for selection signatures with 50K SNPs, a large region under selection on chromosome 6 (33.22 to 41.02 Mbp) was detected and the *ABCG2* (OAR6:36 514 210-36 556 824) and *NCAPG/LCORL* (OAR6:37 256 548-37 333 851/OAR6:37 365 236-37 452 332) were identified as likely candidate genes in this region (Fariello *et al.*, 2014). In our study the same region was detected to be under selection. It was still large (between 22.85 to 48.63 Mbp) however, the most significant single SNPs tests were located in a narrower region of about 2 Mbp (between 35.76 and 37.81 Mbp). We found two clearly distinct haplotypes selected on in this region, one seen only in the Manech Tête Rousse breed and one seen in southern breeds as well as in two northern breeds, the Berrichon du Cher and the Île de France (Figure 1). As shown on our analysis of genetic diversity, both these northern breeds have an history of ancient crossbreeding with a Merino-related population (Figure 4D). When reconstructing a population tree using only SNPs present in this selection signature, these two breeds actually cluster with the Mérinos d’Arles (Figure 1) within the southern group. Therefore we conclude that a selected mutation originating from a southern population related to the Merino has been introgressed and selected again in these two northern breeds, thereby producing by far the strongest signal of selection in our analysis. As this selection event affected a large genome region, it is hard to determine which of the two candidate loci has been targeted by selection. Indeed *ABCG2* is associated with milk production traits in cattle (Olsen *et al.*, 2007) and sheep (García-Fernández *et al.*, 2011; Árnyasi *et al.*, 2013) while *NCAPG* and *LCORL* are associated with growth and height related traits in cattle (Sahana *et al.*, 2015), chickens (Liu *et al.*, 2013), horses (Tetens *et al.*, 2013), humans (Soranzo *et al.*, 2009), pigs (Rubin *et al.*, 2012) and sheep (Al-Mamun *et al.*, 2015; Matika *et al.*, 2016). It is difficult to determine which of the *NCAPG* or *LCORL* gene is causative for these traits because the region exhibits elevated Linkage Disequilibrium in these species. In the Manech Téte Rousse, a hardy dairy breed used for cheese production in the Pyrenees region, **ABCG2** could be considered a better candidate. For the other, introgressed, haplotype, the fact that the region has been under selection in both meat and milk breed could favour the *NCAPG/LCORL* locus as the underlying target.

**Figure 1.**
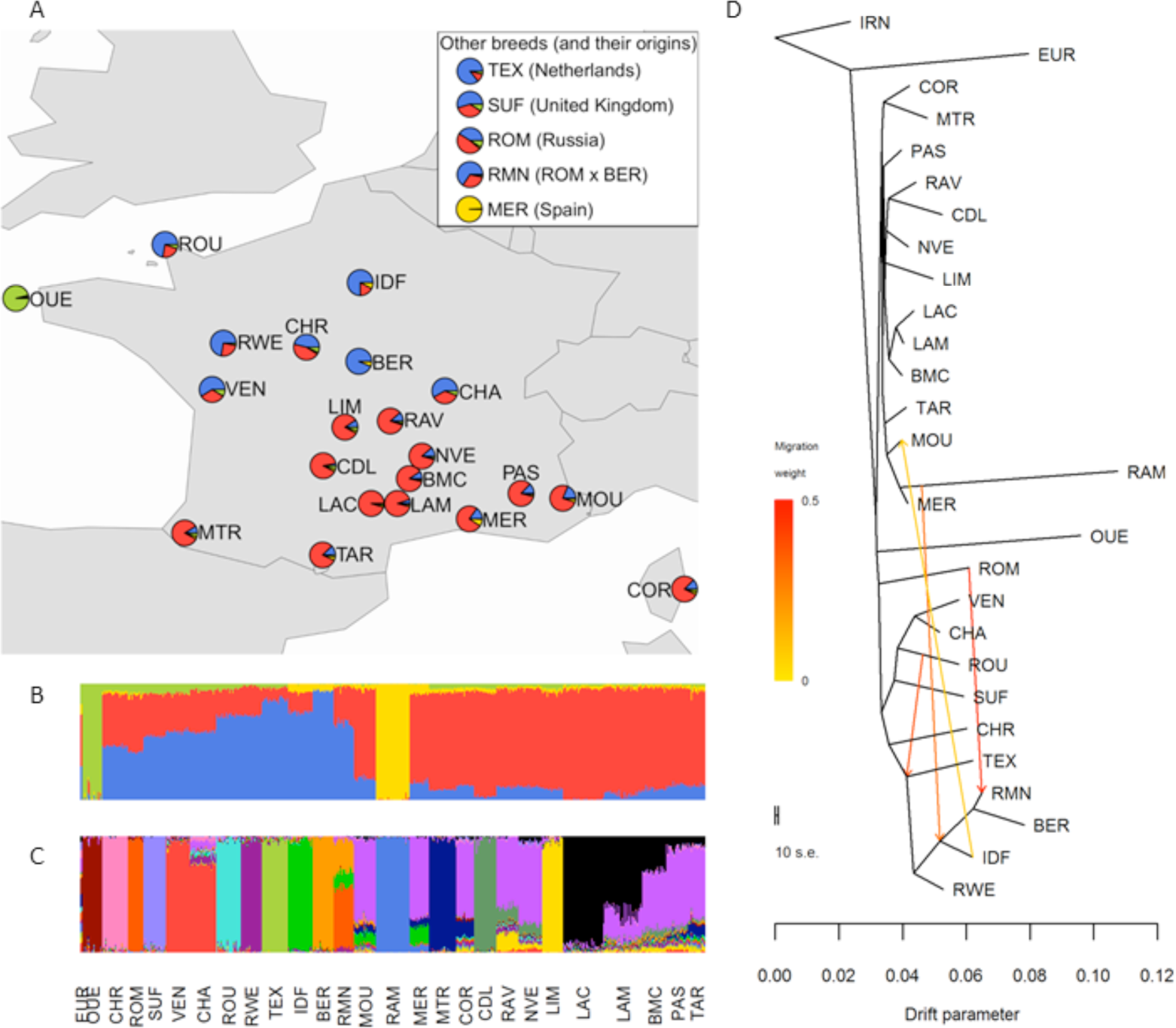
**Population structure of French sheep. (A)** Average individual ancestry coefficients within breed for four ancestral populations. Breeds are mapped to their geographical origin. Individual ancestry coefficients for (B) four ancestral populations and (C) 16 ancestral populations. (D) Maximum likelihood population tree with four migration events

### Confirmation of allelic heterogeneity in MC1R

We chose a selection signature on chromosome 14 (page 36 of Supplementary Table 3), a case of allelic heterogeneity, to investigate further. This signature was chosen because it harbours a gene, *MC1R*, with known causal mutations for black coat colour in sheep and in other mammals (Våge *et al.*, 1999). Coat colour and pattern are a result of pigmentation from melanin. Melanocytes are cells that produce melanin in granules called melanosomes (Birbeck *et al.*, 1956). Eumelanin or pheomelanin is produced depending on whether an agonist, the *melanocyte stimulating hormone* (*α-MSH*), or antagonist, the *agouti signalling protein* (*ASIP*), respectively bind to the *melanocortin 1 receptor* (*MC1R*) (Ducrest *et al.*, 2008). The melanosomes transfer melanin to keratinocytes, the most predominant cell type in the outer layer of skin, the epidermis (Birbeck *et al.*, 1956). Keratinocytes can then incorporate melanin (Forrest *et al.*, 1985). For example, it has been shown in a study of Asiatic sheep that black individuals had only eumelanin present in their wool (Aliev *et al.*, 1990).

Our analysis revealed that selection in this region affected the only three black faced breeds in our dataset: the Romanov, the Suffolk and the Noire du Velay. Furthermore, haplotype diversity plots showed they were clearly selected on different haplotypes (Figure 2A, B). To test whether these three breeds were selected not only on different haplotypes but also on different mutations in *MC1R* we resequenced individuals from these breeds as well as a breed not found to be selected in this region, the Texel.

**Figure 2.**
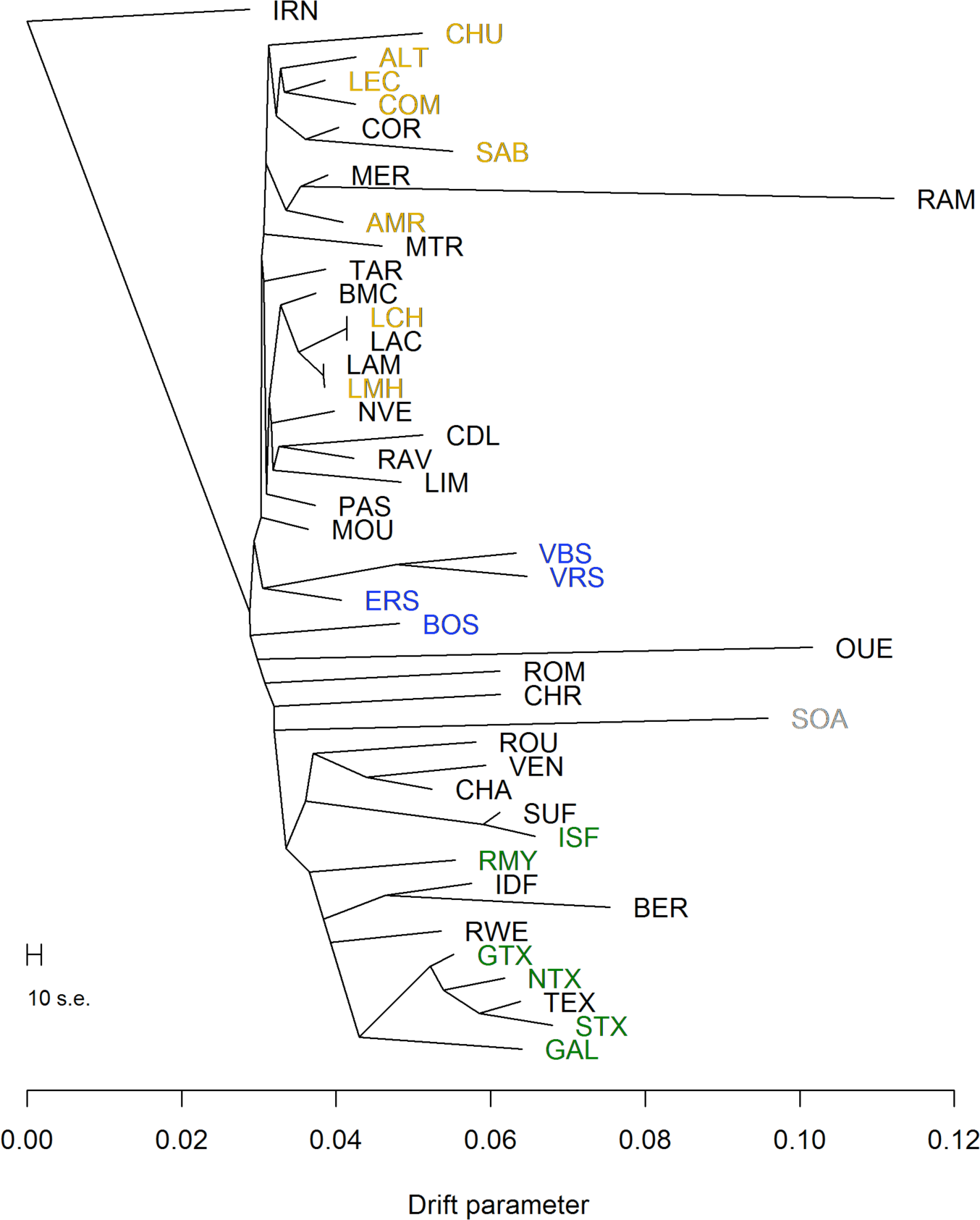
**Maximum likelihood population tree of European sheep breeds estimated using 40K SNPs.** Breeds from this study dataset are in black. SheepHapMap breeds are from Northern Europe ( in grey), Britain (in green), Switzerland (in blue) and West and Middle Mediterranean (in gold).

**Figure 3.**
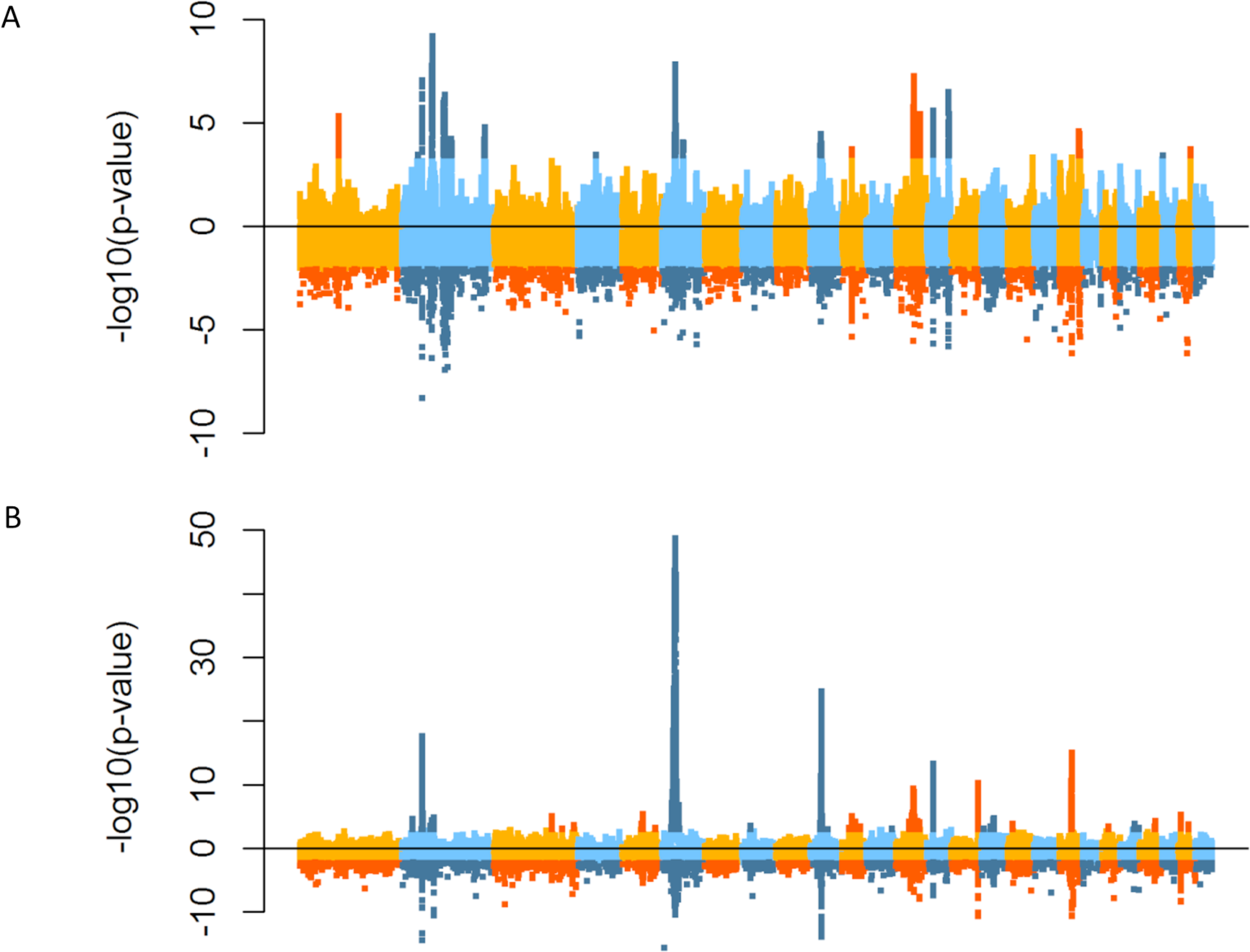
**Manhattan plots of hapFLK and FLK results** in northern (A) and southern (B) populations. In each panel, the positive values are the -log10(p-value) of the hapFLK test and the negative values are the log10(p-value) of the FLK test. Significant SNPs (FDR < 0.05 for hapFLK and < 0.01 for FLK) are shown in darker colors.

**Figure 5.**
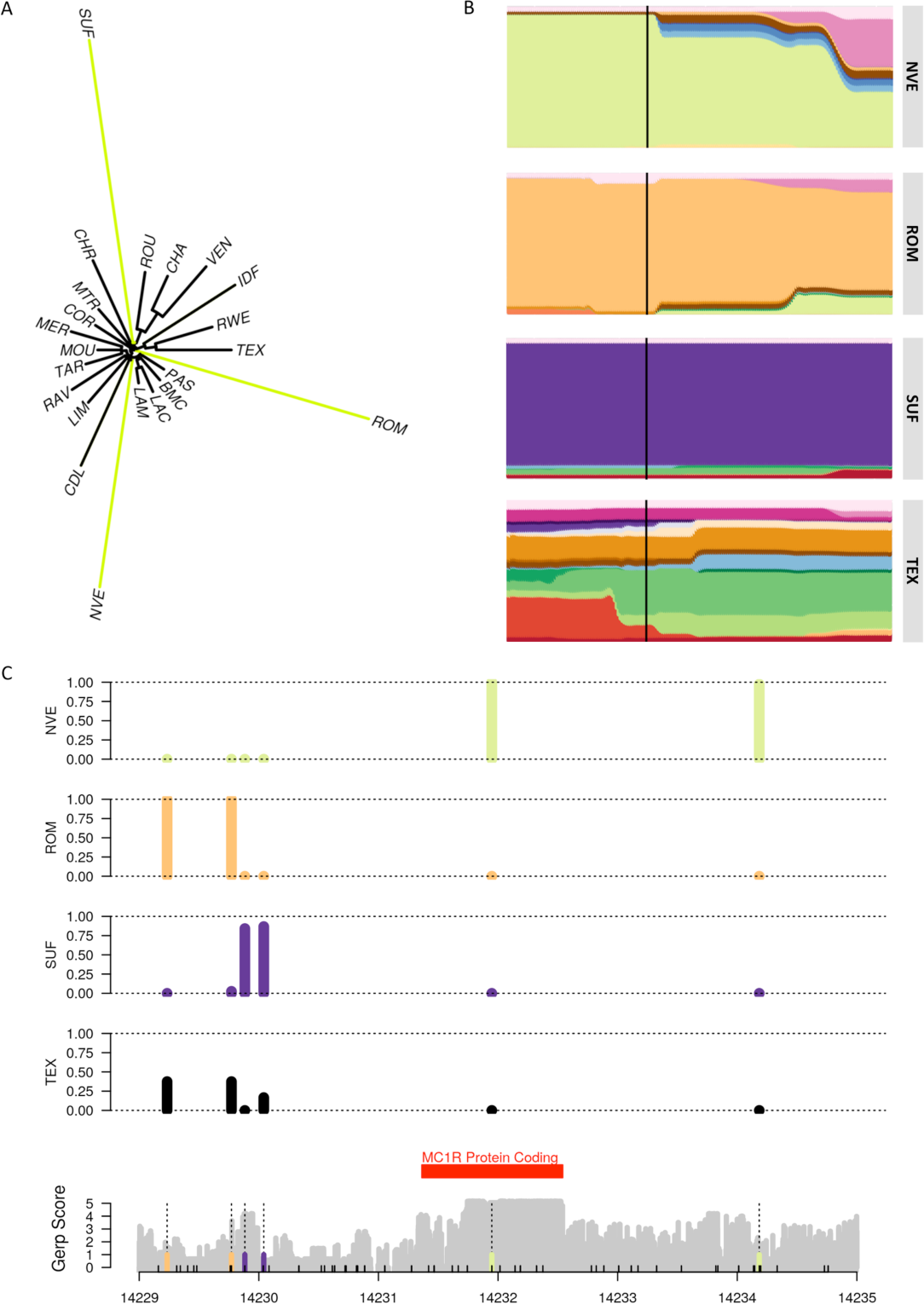
**Chromosome 14:14069040 to 14530851.** (A) Local tree and (B) haplotype cluster frequency plots for Noire du Velay (NVE), Romanov (ROM), Suffolk (SUF) and Texel (TEX) breeds for chromosome 14:14069040 to 14530851 (with the position of MC1R marked with the black line) and (C) mutation location and their frequencies in Noire du Velay (NVE), Romanov (ROM), Suffolk (SUF) and Texel (TEX) breeds

Sequencing of 6 894 bp of the *MC1R* region and genotyping of the Noire du Velay, Romanov, Suffolk and Texel breeds allowed us to generate a reference sequence (EMBL accession number LT594967) with 2 remaining N blocks of 270 and 261 nucleotides in the 5’ region. It exhibits a unique open reading frame, which is consistent with the literature (Fontanesi *et al.*, 2011). We identified a potential TATA box with a “TATAAA” motif ( at positions 14 229 176-14 229 181), 2 522 bp before the start codon and a potential polyadenylation signal “AATAAA” (at positions 14 233 015-14 233 020). Therefore we can assume that the 5’UTR of the sheep *MC1R* transcript is about 2 500 bp long and the 3’UTR around 500 bp long. Few data are present for mammalian 5’UTR of *MC1R* transcripts. The NM_002386.3 human transcript also shows a long 5’UTR (1 380 bp) and a 766 bp 3’UTR. The deduced *MC1R* sheep protein is 317 amino acids, the same number as in cattle (*Bos Taurus*) with a 96% identity between the two proteins.

Our sequences allowed us to detect and further genotype six SNPs and one 11 bp insertion (Figure 2C) (dbSNP accession numbers ss# 1996900605 to 1996900611). We found that the three breeds were indeed almost completely fixed around *MC1R*, and that the three main breed haplotypes were completely different (Supplementary Table 5 and Supplementary Figure 6).

The Noire du Velay resequenced individuals were all homozygotes in the region and carriers of the two known mutations for dominant black in sheep. These mutations (named c.218T > A and c.361G > A in the literature) have been shown to be responsible for dominant black coat in sheep in Norwegian Dala, Damara, Corriedale, Spanish Merino, and Massese breeds (Våge *et al.*, 1999; Våge *et al.*, 2003; Fontanesi *et al.*, 2010).

The Romanov individuals were all homozygotes for the same haplotype in the region, but did not carry the two known mutations above. We identified two mutations that are not found in Noire du Velay or Suffolk. However they were found at intermediate frequency in Texel, a breed where all individuals are white. Although the presence of other coat colour modifier loci in the Texel are possible, it is maybe more likely that the Romanov mutation conferring black coat colour lies outside of the 7 Kb region sequenced here, or in the remaining N blocks of the sequence.

Finally, the Suffolk breed exhibits yet another allele frequency profile different from the two other black headed breeds. We found two mutations of high frequency in the Suffolk, one of which (rs406233740) is a SNP at position 14 229 883 that is not found in any of the other breeds and lies in a region of elevated GERP score, *i.e*. highly conserved among mammals.

Taken together our results confirmed the presence of three different haplotypes around the *MC1R* gene, determined by the combination of seven variants, for Noire du Velay, Romanov and Suffolk breeds. Consistent with the SNP array analysis, the three selected breeds showed almost no between individual variation within breeds, and carried completely different sets of mutations. Assuming *MC1R* is the causative gene, which seems highly likely given the large literature on its effect on coat colour, our results show that only in the Noire du Velay breed coding sequence variants can be the causative mutations. In the two other breeds, regulatory variants are involved, possibly SNP rs406233740 in the Suffolk.

In this study, we presented an analysis of a large set of French sheep breeds based on high density genotyping. We showed that French sheep populations harbour great genetic diversity, with influences from both southern and northern Europe. We detected a large set of selection signatures that we expect will foster new research on studying effects of variation of these genes on phenotypes and shed light into the history of sheep domestication and breeding. In particular, we have showed that some of these regions have been targets of selection on different mutations, in independent genetic pools. This demonstrates the importance of these adaptive regions for agronomical and biological traits in sheep and most likely in other mammalian species. Working with a large set of diverse populations allows to reveal at least partially the history of adaptive mutations and in one particular case through selection on adaptive introgression. Finally, we believe that the combination of this dataset with other likely to come offers great prospects to decipher the history of animal domestication, and its relation to the human neolithic expansion.

## Acknowledgements

CMR benefited from a joint grant from the European Commission and the Swedish University of Agricultural Sciences, within the framework of the Erasmus Mundus joint doctorate “EGS-ABG”. This work was funded by the Animal Genetics division from the French National Institute for Agricultural Research (INRA), the European 7th Framework Programme (3SR project) and the CORAM members (OPA project lead by Jérôme Raoul). The biological samples for the Merinos de Rambouillet were obtained thanks to the CRB-Anim infrastructure, ANR-11-INBS-0003, funded by the French National Research Agency in the frame of the ‘Investing for the Future’ program. The funders had no role in study design, and analysis, decision to publish, or preparation of the manuscript. We deeply appreciated the involvement of various breeding organizations for their assistance in sample collection (France Génétique Elevage http://en.france-genetique-elevage.org/, FranceAgriMer http://www.franceagrimer.fr/, le Groupement des Moutons d’Ouessant http://www.moutons-ouessant.com/ and l’Institut de l’Élevage (http://idele.fr), especially Coralie Danchin as coordinator of the GenImpact program. Finally, we thank Julie Demars for the European mouflon samples, François Pompanon for providing the Asian Mouflon allele frequencies and LABOGENA (http://www.labogena.fr/), especially Julie Ogereau and Bérengère Camus, for the genotyping using Illumina Ovine Infinium^®^ HD SNP Beadchips.

